# Phenological stage influences how white lupin (*Lupinus albus*) root exudate metabolites respond to phosphorus supply

**DOI:** 10.64898/2026.05.28.728569

**Authors:** Sasha Pollet, Jean-Thomas Cornelis, Thorsten Knipfer, Cindy Prescott, Kylee Tate, Young-Mo Kim, Guillaume Lobet

## Abstract

The composition and quantity of root exudates are strongly influenced by physiological and environmental conditions, reflecting dynamic changes in plant metabolism. Although many studies report that root exudate metabolite profiles vary with plant phenology, few have disentangled the effects of phenological stage from those of nutrient availability. We collected root exudates from white lupin *(Lupinus albus*) grown in a fine phosphorus (P) gradient (5, 10, 20, 30 and 50 µM P) in hydroponics at three developmental stages, performed untargeted metabolomics using GC-MS, and measured 10 above and belowground traits. Our results show that plant phenological stage exerts a stronger influence on exudate metabolomic profiles than variation in P supply. During leaf development, exudates were dominated by metabolites associated with carbon metabolism, whereas flowering was characterized by compounds related to secondary metabolism and cell wall turnover. Phosphorus influenced exudate profiles only at the flowering stage, with distinct profiles observed at 5 and 10 µM P compared with 20-50 µM P. These findings provide new insights into the temporal regulation of root exudation and demonstrate that plant developmental stage is a primary determinant of metabolic responses to phosphorus availability and, potentially, rhizosphere functioning under nutrient limitation.

**Highlight:** Root exudate quantity and composition in white lupin is shaped more by plant phenological stage than phosphorus supply, with phosphorus effects emerging only during flowering.

## Introduction

Root exudates are soluble organic compounds released by living roots into the soil comprising high (>1000 Da; e.g. enzymes, mucilage) and low (MW;<1000 Da; e.g. sugars, organic acids, phenolics) molecular weight compounds. Chemically very diverse, root exudates encompass both primary (metabolites that are essential for plant growth and functioning e.g. organic acids, carbohydrates, amino acids) and secondary metabolites (not involved directly in cell growth and maintenance e.g. phenolics coumarins, phytosiderophores). Together, these compounds can represent a significant proportion of plant photosynthates released in the soil, up to 25% of recently assimilated C (Lynch and Whipps, 1990; Badri *et al*., 2008; Pausch and Kuzyakov, 2018).

Root exudate metabolite profiles, defined as the detailed composition and concentration of various metabolites present within a sample (Liu *et al*., 2025), are highly variable and reflect underlying plant metabolic activity (Canarini *et al*., 2019; McLaughlin *et al*., 2023). Most of the exudates originate from non-structural carbohydrates produced in the leaves through photosynthesis. The fate of carbohydrates and their movement within plants is highly dynamic and shaped by environmental conditions and developmental stage (Hartmann *et al*., 2020). Their transport depends on source-sink activities (Münch, 1930). During vegetative growth, a large proportion of assimilated C supports root and shoot expansion, whereas reproductive development redirects C towards flowers, fruits, and seeds, substantially altering internal C partitioning and shoot–soil C fluxes (Hartmann *et al*., 2020; Ren *et al*., 2024). Consistent with this, changes in exudate profile over phenological stages have been highlighted for different plant species (Ritter *et al*., 2025) : in maize, (Zea mays; Santangeli et al., 2024), wild oats (Avena spp.; Zhalnina et al., 2018), cucumber (Cucumis sativus; Feng et al., 2023), and thale cress (Arabidopsis thaliana; Chaparro et al., 2013).

Besides plant phenological stage, exudate metabolite profiles depend on multiple factors, including plant species and genotypes (Robert *et al*., 2026), environmental conditions such as drought, temperature and elevated CO_2_ (Wasaki *et al*., 2005; O’Sullivan *et al*., 2021), nutrient availability and stoichiometry (Carvalhais *et al*., 2011), and interaction with rhizosphere microbes (Ma *et al*., 2022). Under suboptimal conditions such as drought or nutrient limitation, plant growth becomes decoupled from C assimilation: leaf expansion is restricted earlier than photosynthesis, leading to an accumulation of carbohydrates in source tissues (White and Hammond, 2016). This surplus C is subsequently redistributed to sink organs and may be exported belowground where it can support symbiotic microorganisms or be exuded from roots (Prescott *et al*., 2020; Bunn *et al*., 2024; McCulloch *et al*., 2026). Capitalizing on this surplus C exudation under optimized suboptimal conditions is suggested as a strategy to increase soil organic matter content and enhance nutrient acquisition in cropping systems without substantially compromising aboveground biomass (Prescott *et al*., 2021; Wang *et al*., 2025).

Among nutrients, phosphorus (P) has received particular attention for its strong influence on root exudation (Hinsinger, 2001; Jones *et al*., 2004). It has been shown that reducing P fertilization by up to 50% of the conventional recommended dose increased root exudation by 28%, while only decreasing plant biomass by 2.2% (Wang *et al*., 2025). Phosphorus limitation commonly induces increased exudation of protons, acid-phosphatase, low-molecular-weight organic acids, particularly carboxylates, which enhance P solubilization and mobilization in soils (Neumann and Römheld, 2011). Alongside nitrogen (N), P is one of the most limiting nutrients for plant growth worldwide (Vitousek *et al*., 2010). Plant acclimation to P limitation involves a suite of metabolic, physiological, and morphological adjustments aimed at maintaining essential cellular functions (White and Hammond, 2016). For example, cell membrane phospholipids can be replaced by P deprived components like sulfo- and galactolipids and metabolic bypass reactions can occur, such as modified glycolysis (Plaxton and Tran, 2011). Changes in metabolic pathways in response to plant development or environmental conditions can result in altered exudation patterns (Lynch and Brown, 2016).

To date, the effects of different P supply on root exudation have been investigated primarily at specific developmental stages, among different genotypes within a single plant species or across species (Vengavasi and Pandey, 2018; Pantigoso *et al*., 2020a). Although several studies report that both the rate and composition of root exudation vary with plant phenology, only one study has explicitly disentangled the effect of developmental stage from those of nutrients’ availability. Using *Arabidopsis thaliana*, Pantigoso et al., 2020 showed that P supply shapes root exudate profiles differently at the vegetative and, more markedly at the bolting stage, with increased phosphate availability leading to reduced exudation of metabolites involved in P solubilization.

Untargeted metabolomics is a powerful approach to capture the diversity of root exudates and to investigate how exudate profiles respond to environmental and physiological changes (Salem et al., 2022; Brown et al., 2024). While specific compound classes involved in nutrient acquisition or plant–microbe interactions have been extensively studied, the role and origin of most exuded metabolites remain uncharacterized and likely reflect broader metabolic and stoichiometric adjustments to internal and external constraints (McLaughlin *et al*., 2023). These complex exudate cocktails interact with microorganisms and minerals in the rhizosphere, thereby influencing soil biogeochemistry and ultimately impacting plant performance and resilience in agroecosystems (Badri and Vivanco, 2009; Williams and de Vries, 2020; Oburger *et al*., 2022; Han *et al*., 2023).

In a previous study, we showed that exudate cocktails produced by white lupin (*Lupinus albus*) in a P gradient at a specific growth stage had strong effects on mineral dissolution depending on soil developmental stage (Pollet *et al*., 2026). However, because root exudation is a dynamic process, exudate profiles might change across phenological stages and respond differently to P supply, ultimately altering exudate fate in the rhizosphere. White lupin is well known for exuding organic acids, particularly citrate and malate, from cluster roots under P limiting conditions. However, while Müller et al., 2015 compared untargeted metabolomic profiles of shoot, root and cluster from P deficient plant and P sufficient plants at an early developmental stage (15 days), no study, to our knowledge, has examined white lupin exudates metabolomic profile across multiple phenological stages under varying P supply.

Here, we used a gas chromatography-mass spectrometry (GC–MS)-based metabolomic profiling approach to characterize temporal changes in white lupin root exudate metabolite profiles under a fine gradient of P supply (5, 10, 20, 30 and 50 µM P) in a hydroponic system. The main objective of this study was to characterize changes in exudate profiles depending on P supply and phenological stages. Because of the high exudation rates of white lupin cluster roots, we hypothesize that root exudate profiles will reflect cluster root activity and will change dynamically over the course of plant development and in response to P supply.

## Material and Methods

### Plant cultivation

We cultivated white lupin (*Lupinus albus* L. cv. Kiev) following the protocol described in Pollet et al., 2026. Briefly, we grew plants hydroponically in a controlled growth chamber for seven weeks in fall 2023. Plants were grown in a gradient of P concentrations (5, 10, 20, 30, or 50 µM P), with P supplied in the form of KH□PO□, while all other nutrients were provided at non-limiting concentrations. After five days of germination in a solution containing 1 mM CaCl□ and 5 µM H□BO□ (Shane *et al*., 2003; O’Sullivan *et al*., 2021), we transferred the seedlings to 25-L black containers containing half strength nutrient solution. Once two fully expanded leaves had developed, we transferred plants to the final treatment solutions. We adjusted nutrient solution pH to 6 and renewed the solution every five days. Growth conditions consisted of a 16 h light/8 h dark photoperiod, day/night temperatures of 24/18 °C, 50% relative humidity, and a light intensity of 400 µmol m□² s□¹. We removed cotyledons at developmental stage 1.2 to induce early P deficiency.

The first exudate collection happened three weeks after plant emergence, when seven leaves had emerged from the bud, using five plants per P treatment. Four hours after the onset of light, we rinsed root systems three times in distilled water and transferred them into 150 mL foil-wrapped beakers containing 130 mL distilled water for a two-hours exudate collection period (Oburger and Jones, 2018). During exudate collection, we measured photosynthesis on three plants per P treatment from the last fully developed leaf using a LI-COR 6800.

Following collection, we filtered exudate solutions at 0.5 µm, freeze-dried and stored them at −20 °C prior to metabolomic analysis. We then harvested shoots, roots and measured total leaf area using a LI-3000A leaf area meter. We scanned root systems from three or four replicates per treatment in a transparent water tray using an Epson V850 Pro flatbed scanner. We quantified total root length and root tip number (excluding cluster root tips) using *RhizoVision* software (Seethepalli and York, 2020), while we manually delineated cluster roots using *SmartRoot* (Lobet *et al*., 2011). We dried root and shoot tissues at 60 °C for 24 h before weighting them. The same procedure was repeated five and seven weeks after emergence, corresponding to the stem elongation and flowering stages, respectively (**Table 1**).

**Table 1:** Sampling time and corresponding phenological stage described in New South Wales. Department of Industry and Investment, 2011.

| Sampling time (week after emergence) | Phenological stage |
| --- | --- |
| Week 3 | 1.7 (leaf emergence- 7 leaves emerged from the bud) |
| Week 5 | 2.5 (stem elongation - Bases of several leaves clearly separated from each other) |
| Week 7 | 3.2 (flowering - Hooded bud stage) |

### Metabolomics analysis

For the metabolomics analysis, we reconstituted the lyophilized material in 2 mL nanopure water. We used 1 mL for gas GC-MS analysis and retained the remaining 1 mL as backup. We vortexed the reconstituted sample in an Eppendorf tube before transferring them to glass vials and drying them completely using a speed vacuum concentrator. Once dried, we chemically derivatized the samples and analyzed them by GC-MS following the protocol described by (Xu *et al*., 2018). We performed global metabolomic profiling by comparing GC-MS spectra against our in-house expanded version of the Agilent Fiehn metabolite database, utilizing two-dimensional parameters such as fragmented mass spectra and retention indices (Kind *et al*., 2009). We further confirmed metabolite identities using the NIST23 and Wiley (11th Edition) databases. All the metabolite identifications reported here are based on the level 1 identification recommended by Metabolomics Standards Initiatives (MSI) (Sumner *et al*., 2007). We classified metabolites into chemical groups and subgroups using PubChem (Kim *et al*., 2025) and KEGG (Kanehisa *et al*., 2025) databases.

For absolute quantification of selected metabolites, we generated calibration curves using serial dilutions of standard compounds. We prepared standard references for aspartic acid, citric acid, fructose, glucose, glutamic acid, malic acid, malonic acid, oxalic acid, succinic acid, and sucrose in a 50:50 methanol:water mixture. We then diluted the standard to final concentrations of 0, 0.1, 1, 10, and 100 µg/mL. We analyzed triplicate standards with duplicate technical injections by GC–MS after sample analysis. All calibration curves produced coefficients of determination (R²) greater than 0.95.

### Statistical analysis

We used two-ways analysis of variance (ANOVA) to assess the effect of P treatment and phenological stage on plant traits and root exudates (n=5) (R Core Team, 2022). We tested residual normality with Shapiro–Wilk and Q–Q plots and homogeneity of variance using Levene’s. We performed Tukey’s HSD post hoc test to identify significant differences between group means. We log- or square-root transformed the data when assumptions were not met. If assumptions could not be met despite transformation, we used Welch’s ANOVA and the Games–Howell post hoc test (via the rstatix package (Kassambara, 2023)).

For exudate data analysis, we expressed results both on a per-plant basis and standardized by root biomass. For root biomass standardization, we divided the absolute concentration or GC–MS peak area of each detected compound by the corresponding root biomass. For quantified compounds (absolute concentrations), we performed a two-way ANOVA to test the effects of P treatment, phenological stage, and their interaction on compound exudation.

For the untargeted global metabolomics analysis, we used 5% thresholds for peak area data filtering, both for low-variance and low-abundance filtering. We then log-transformed and mean-centered the data. We conducted multivariate analyses using MetaboAnalyst, a web-based metabolomics data processing tool (http://www.metaboanalyst.ca) (Pang *et al*., 2024). First considering the entire dataset, we performed a principal component analysis (PCA) to visualise the data and use a permutational multivariate analysis of variance (PERMANOVA) to test whether the centroids of the different groups (P supply and phenological stage) were significantly different from one another. Because we did not observe strong effect of P supply across all phenological stages, we subsequently assessed the effect of P supply separately within each phenological stage by performing a PCA, a PERMANOVA and one-way ANOVA followed by a Tukey HSD test for each metabolite to test for P treatment effects. We also performed hierarchical clustering heatmaps by using Euclidean distance and the Ward clustering method (Dietz *et al*., 2020). To evaluate the effect of phenological stage on the exudate profile independently of P treatment, we performed a PCA, PERMANOVA and hierarchical clustering heatmaps on the full dataset without accounting for P treatment. We used a one-way ANOVA followed by a Post Hoc Tukey HSD test on each metabolite to highlight significant difference in means across the phenological stages. As described above, we conducted all analyses for both peak area both expressed on a per-plant basis and standardized by root biomass.

## Results

### Above and belowground plant traits response to varying P supply across phenological stages

Three weeks after emergence, plants had produced seven leaves from the bud. By week five, stem elongation was evident, as the base of several leaves was clearly separated. By seven weeks, plants reached the hooded bud stage of flowering, marking the transition to the reproductive phase. Overall, the effect of P on aboveground traits dependent on the plant phenological stage, whereas belowground traits were influenced by P treatment and phenological stage separately (*Supplementary material 1*).

Root system architecture and root biomass were influenced by P treatment and phenological stage but not by their interaction. Specific root length (SRL), number of root tips (excluding cluster rootlet), total root length, number of cluster roots, and total length of cluster were all affected by plant phenological stage. Specific root length significantly decreased from leaf development to flowering stage (**Fig. 1D**). In contrast, the number of root tips (excluding cluster roots), the number of cluster roots, and total root length increased from leaf development to stem elongation stage but did not increase further before the reproductive phase. Root biomass, however, continued to increase until flowering. Phosphorus treatment significantly affected root biomass, the number of cluster roots, and the proportion of total root length occupied by cluster roots. Root biomass was lowest under the 5 µM P treatment and did not differ from 50 µM P, which also did not differ from 10, 20, or 30 µM P. The number of cluster roots under 50 µM P was significantly lower than under 10 µM P, while the other P treatments did not differ from either. In contrast, the proportion of total root length occupied by cluster roots increased as P supply decreased (**Fig. 1C**).

**Fig. 1:**
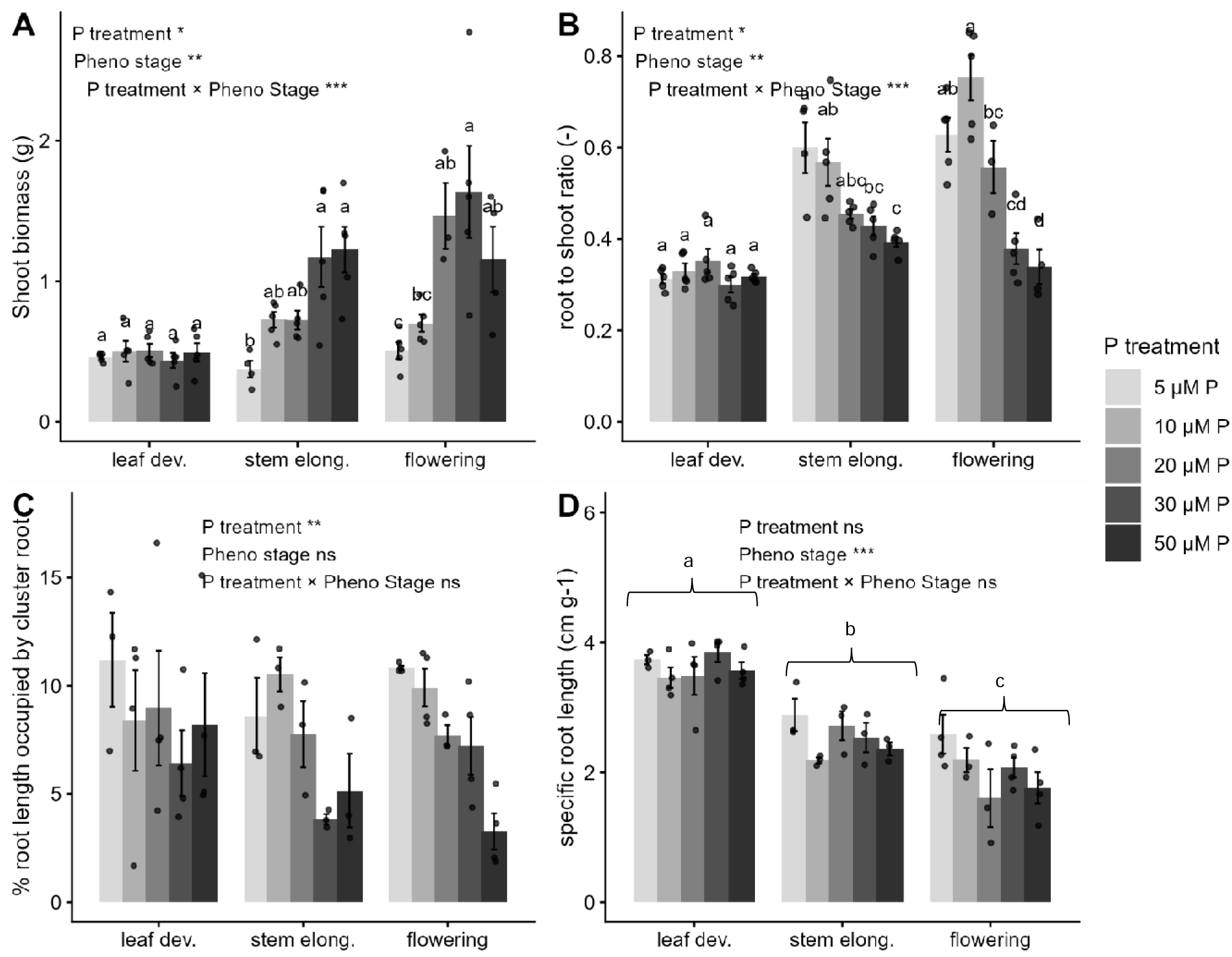
Plant traits across P supply and phenological stages. (A) Shoot biomass, (B) root-to-shoot ratio, (C) percentage of total root length occupied by cluster roots and (D) specific root length. Data are presented as means ± SE, with each dot representing an individual observation. Effects of phenological stage, P treatment, and their interaction were assessed using two-way ANOVA. Significance levels are indicated as follows: P < 0.05 (*), P < 0.01 (**), P < 0.001 (***), and not significant (ns, P > 0.05). Where significant interactions occurred, different letters indicate significant differences among P treatments within each phenological stage (Tukey’s HSD test, P < 0.05). Different letters on top of brackets indicate significant differences among phenological stages.

Shoot biomass, root-to-shoot weight ratio (R:S), leaf area, and specific leaf area (SLA) showed significant interactions between phenological stage and P supply, indicating that P effects varied across phenological stages. During leaf development, none of the measured shoot or root parameters responded to P availability. During stem elongation, P effects became significant for shoot biomass, R:S, leaf area, and SLA. Shoot biomass and leaf area were significantly lower under 5 µM P than under 30 and 50 µM P. At flowering stage, P supply had a clear effect on shoot biomass, R:S, leaf area, and SLA. Shoot biomass under 20, 30, and 50 µM P was higher than under 10 and 5 µM P but did not differ among these higher P treatments (**Fig. 1A**). Root-to-shoot ratio peaked at 10 µM P and decreased with increasing P supply (**Fig. 1B**). Specific leaf area was the lowest at 5 µM P. The impact of P on photosynthesis started during stem elongation and showed a distinction between the low (5-10 µM P) and med-high P treatment (20, 30 and 50 µM P).

### Higher exudation rate of quantified organic acids, amino acids and sugars at early developmental stage

We quantified seven organics acids (citric, malic, fumaric, succinic, oxalic, tartaric and malonic acids), three sugars (sucrose, fructose and glucose) and two amino acids (L-glutamic and L-aspartic acids). Globally, the exudation rate of these compounds per g of root decreased as plant aged, with some interaction between P treatment and phenological stage for some compounds (*Supplementary material 2*).

First, succinic and tartaric acids, as well as sugars (fructose, sucrose and glucose- **Fig. 2C**) were only dependent on plant phenological stage, and there were highest during leaf development and declined thereafter, with no further differences between stem elongation and flowering. Across phenological stages, tartaric acid responded consistently to P supply, showing higher concentration at 5 µM P than at 30 and 50 µM P, while 10 and 20 µM P were not significantly different from either extreme.

**Fig. 2:**
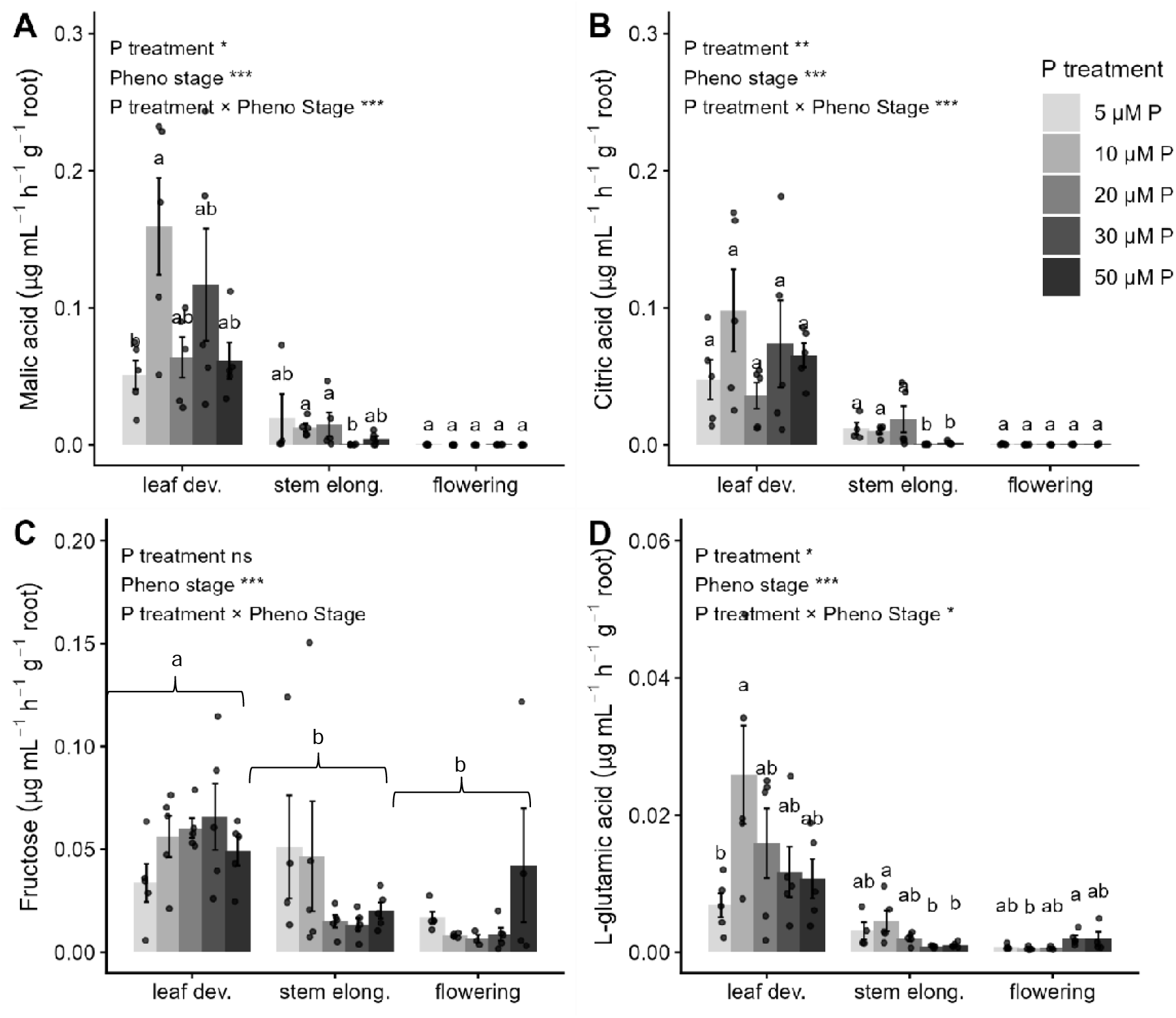
Most abundant quantified metabolites across P treatments and phenological stages. (A) malic acid, (B) citric acid, (C) fructose and (D) L-glutamic acid. Data are presented as means ± SE, with each dot representing an individual observation. Effects of phenological stage, P treatment, and their interaction were assessed using two-way ANOVA. Significance levels are indicated as follows: P < 0.05 (*), P < 0.01 (**), P < 0.001 (***), and not significant (ns, P > 0.05). Where significant interactions occurred, different letters indicate significant differences among P treatments within each phenological stage (Tukey’s HSD test, P < 0.05). Different letters on top of brackets indicate significant differences among phenological stages.

The effect of P on the rest of quantified compounds varied with phenological stage. During leaf development, the concentration of malic (**Fig. 2A**), oxalic acid as well as L-aspartic acids were the highest at 10 µM P. During stem elongation, citric acid exudation was significantly lower at high P (30 and 50 µM) than at low to moderate P (5, 10, and 20 µM) (**Fig. 2B**). Fumaric and malic acids followed a similar pattern, with the lowest concentrations observed at 30 µM P. Amino acids exudation showed an opposite trend, with both L-glutamic and L-aspartic acids significantly higher in 10 µM P than in 30 and 50 µM P (**Fig. 2D**). During flowering stage, only L-glutamic acid was impacted by P treatment, being significantly higher at 30 µM P than in 10 µM P. We quantified malonic acid in some root exudate samples, but not consistently across phenological stages and P treatments.

All quantified organic acids, amino acids, and sugars, except glucose and oxalic acid, showed the same trend when normalized to root biomass or expressed on a per-plant basis; all higher during leaf development than stem elongation and flowering. Oxalic acid exudation per plant basis was constant across P treatment and phenological stages.

### Untargeted metabolomics of white lupin root exudates

We identified a total of 109 compounds in white lupin root exudates by GC-MS, including 28 organic acids, 17 carbohydrates, 11 polyols, 21 amino acids, 9 phenolics, 11 fatty acids and lipid-related compounds, 3 nucleotides/bases, 2 amines, 2 other nitrogen-containing compounds, 1 vitamin, 3 inorganic compounds, and 1 xenobiotic compound (full list of compounds in *Supplementary material 3*).

Overall, phenological stage had a much stronger effect on the root exudate metabolite profile than P treatment (PERMANOVA, p < 0.001 for phenological stage, not significant for P), regardless of whether peak area was normalized to root biomass or expressed on a per-plant basis.

#### Root exudate metabolite profiles across phenological stages

When peak areas were normalized to root biomass (root biomass–standardized peak area), 81 compounds (listed on the heatmap **fig. 4A**) differed significantly across phenological stages. For most of these compounds (60 out of 81), peak areas were higher during the leaf developmental stage than at later stages. No compounds had a significantly higher root biomass-standardized peak area during flowering.

In addition to the higher peak area at early stages, multivariate analysis revealed a clear shift in exudate profile from leaf development to flowering (p = 0.001). This shift is reflected by the separation of sample ellipses along PC1 in the PCA (**fig. 3A**). Compounds with positive loadings on PC1 were predominantly organic acids (citric acid, malic acid, and fumaric acid), amino acids (L-glutamic acid, L-ornithine, L-pyroglutamic acid, and L-tyrosine), and the glucose. In contrast, negative loadings on PC1 were mainly associated with more complex carbohydrates (cellobiose, 1,4-D-xylobiose, and xylose), the polyamine putrescine, lipid-related compounds (adipic acid, caprylic acid, and capric acid), and the aromatic acid 2-furoic acid.

**Fig 3:**
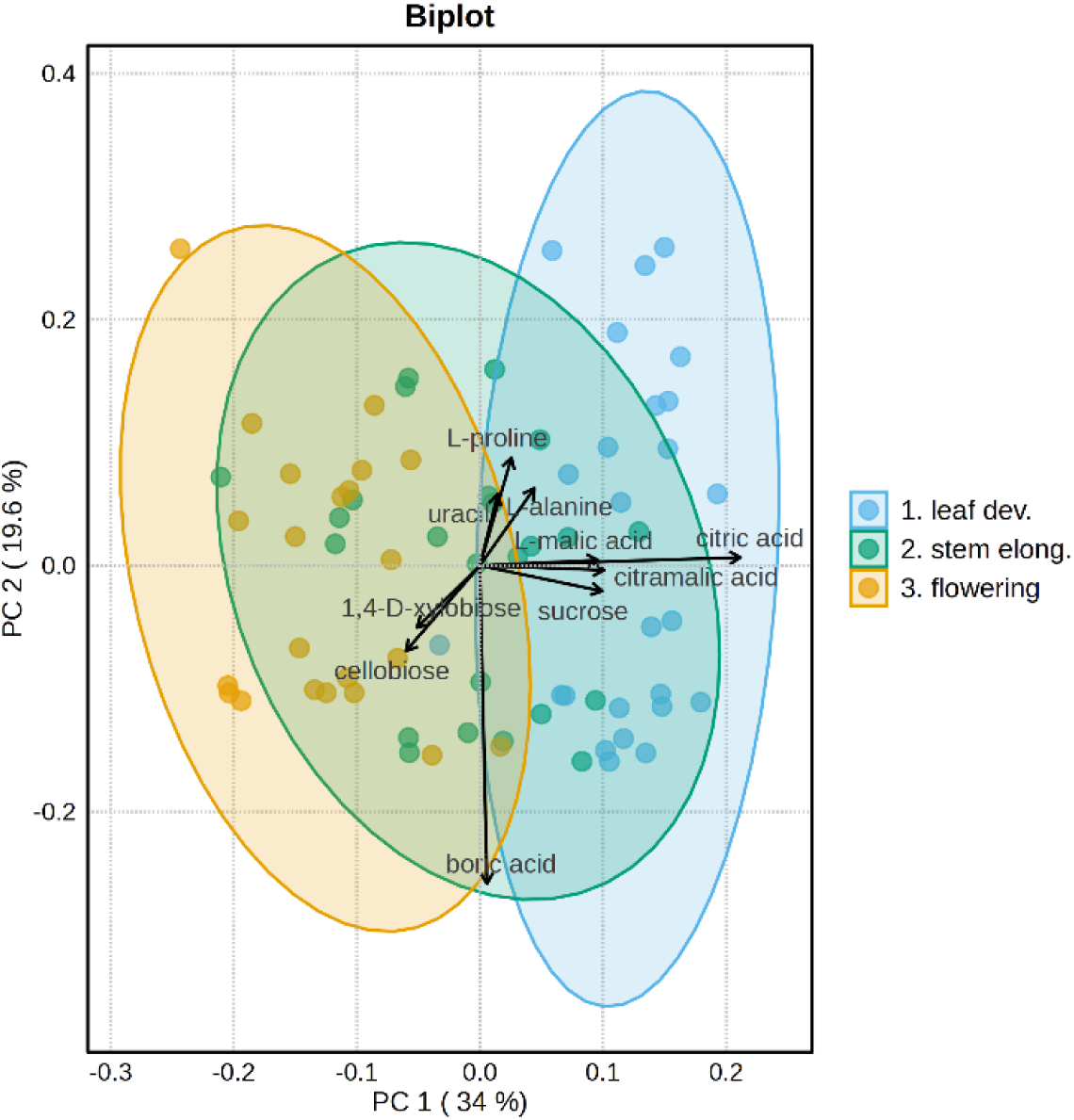
Principal component analysis on the exudate profiles across phenological stages based on peak areas standardized by root biomass, log-transformed and mean-centered prior to analysis. Phenological stage significantly affected exudate profiles (PERMANOVA: F-value: 34.02; R-squared: 0.48935; p-value (based on 999 permutations): 0.001). The 10 features contributing most strongly to the PCA are shown.

When peak areas were expressed on a per plant basis (i.e. total peak area in the exudates solution, not standardized by root biomass), 76 compounds were significantly different across the phenological stages (**fig 4. B** heatmap). Only 24 out of 76 compounds were higher at leaf development compared with flowering. The remaining compounds showed peak area distributed across stem elongation and flowering. Nevertheless, multivariate analysis showed that these exudate profiles were highly similar to those obtained after standardization by root biomass (*Supplementary material 4*).

**Fig 4:**
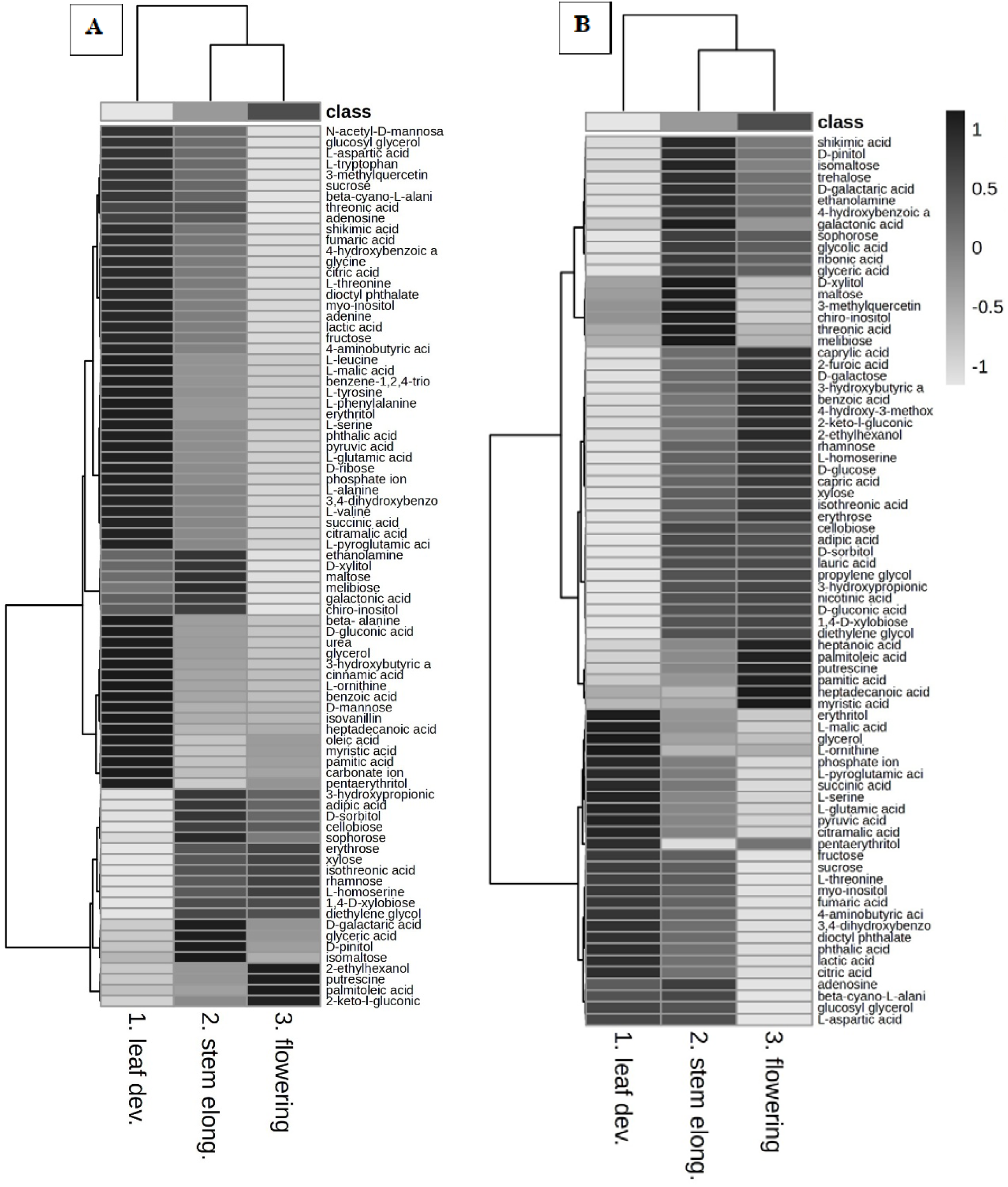
Heatmaps of metabolites showing significant differences among phenological stages (one-way ANOVA, P < 0.05). Hierarchical clustering was performed using Euclidean distance and Ward’s method. Darker shading indicates higher peak areas, whereas lighter shading indicates lower peak areas. (A) Peak areas standardized by root biomass. (B) Peak areas expressed on a per-plant basis.

Overall, these results indicate that, for most compounds, the amount exuded per gram of root was highest during early development (leaf development, and to some extent stem elongation). However, when expressed on a per-plant basis, exudation was more evenly distributed across phenological stages, highlighting the influence of root biomass on total measured metabolites.

#### Interactive effect of P treatments and phenological stages on root exudate metabolite profiles

Across the three phenological stages, no compounds showed changes in root biomass–standardized peak area that were only attributable to the P treatment. When peak area was expressed on a per-plant basis, four compounds (chiro-inositol, D-xylitol, D-sorbitol, and lactic acid) were significantly affected by P treatment, irrespective of phenological stage. Peak area of D-xylitol and D-sorbitol were higher at high P (30 and 50 µM P), whereas lactic acid was higher at 10 and 20 µM P. Chiro-inositol was lower at 5 µM P. The effect of P on the peak area of 28 compounds was dependent on phenological stage.

How P treatment affected exudates during leaf development and stem elongation was similar between root-biomass standardized peak area and expressed on a per-plant basis. During leaf development stage, P had no detectable effect on the root exudate metabolite profile. Similarly, during stem elongation stage, the overall exudate profile did not differ significantly among the P treatments. However, several individual compounds showed significant difference in peak area between P levels (p>0.05), including citric acid, fumaric acid, lactic acid, putrescine, beta-cyano-L-alanine. For all these compounds, peak areas tended to be higher under low to moderate P treatment (05, 10 and 20 µM P) compared with moderate to high P supply (30 and 50 µM P).

At the flowering stage, PCA revealed a significant difference (p = 0.032) and separation between low P treatments (5 and 10 µM P) and moderate to high P treatments (20, 30, and 50 µM P). A total of 21 compounds (root biomass standardized peak area) and 35 compounds (per-plant basis) differed between P levels (**Fig. 6 A. B.**).

For peak areas standardized to root biomass, samples from low P treatments clustered in the quadrant defined by positive PC1 and negative PC2 loadings (**Fig. 5**). In contrast, samples from moderate to high P treatments were largely absent from this region and instead distributed along the positive side of PC2, with little separation along PC1. Positive loadings on PC1 were primarily associated with the following compounds: cellobiose, sucrose, glucosyl glycerol and D-galactaric acid, while negative PC1 loadings with L-alanine, 3-aminoisobutyric acid, uracil, L-leucine and L-valine. Positive loadings on PC2 were characterized by D-xylitol, 3-aminoisobutyric acid, myo-inositol, L-tryptophan, D-sorbitol, and negative loadings by cellobiose, glyceric acid, 2-ethylhexanol, lactic acid, uracil.

**Fig 5:**
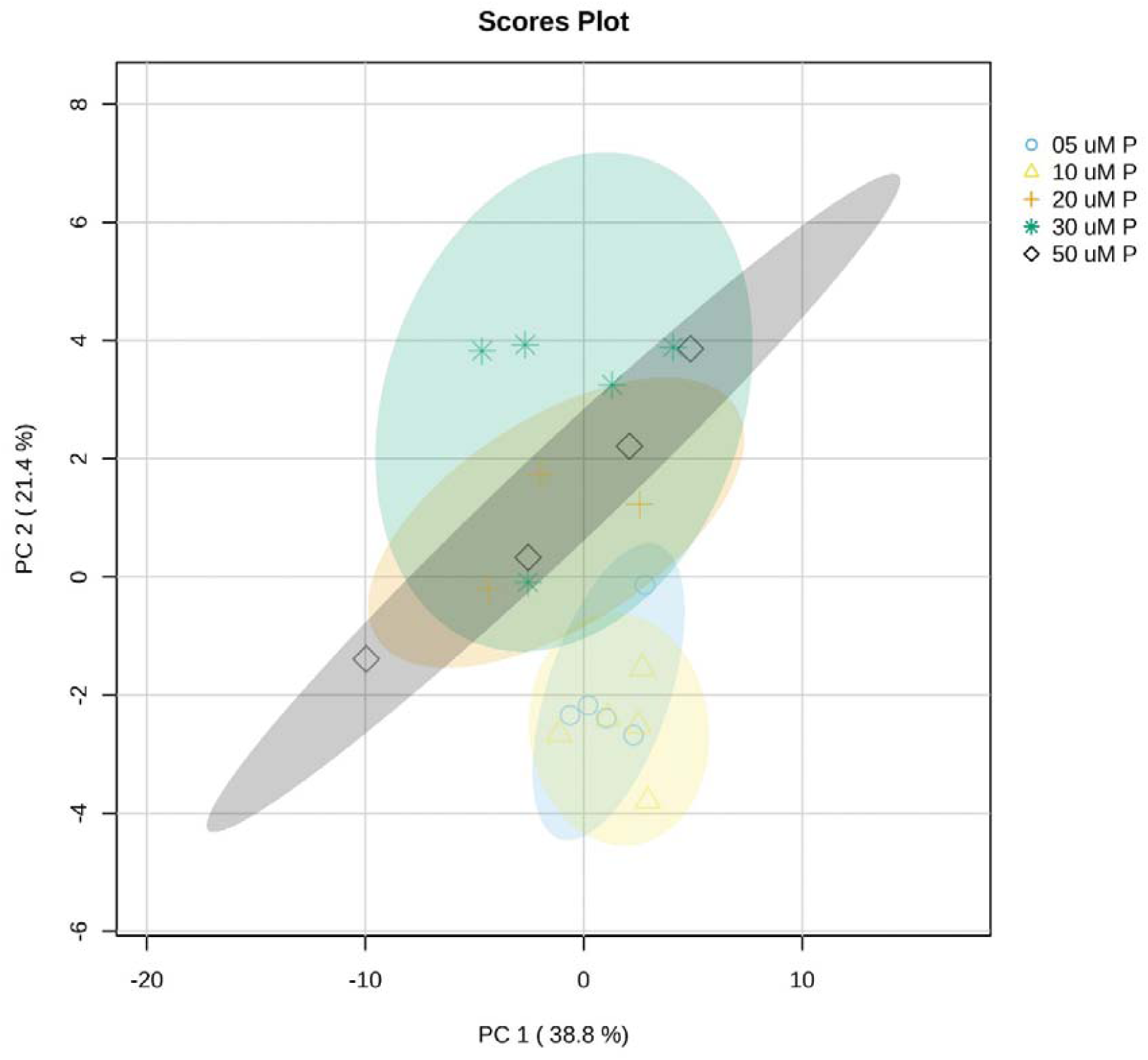
Principal component analysis on the exudate profiles across P treatments at flowering, based on peak areas standardized by root biomass, log-transformed and mean-centered prior to analysis. Phosphorus treatment significantly affected exudate profiles at flowering (PERMANOVA: F-value: 2.4026; R-squared: 0.36116; p-value (based on 999 permutations): 0.04). The 5 features contributing the most to PC1 positive loadings are cellobiose, sucrose, glucosyl glycerol, D-galactaric acid. PC1 negative loadings: L-alanine, 3-aminoisobutyric acid, uracil, L-leucine, L-valine. PC2 positive loadings: D-xylitol, 3-aminoisobutyric acid, myo-inositol, L-tryptophan, D-sorbitol. PC2 negative loadings: cellobiose, glyceric acid, 2-ethylhexanol, lactic acid, uracil. Adapted from Pollet et al., 2026.

**Fig 6:**
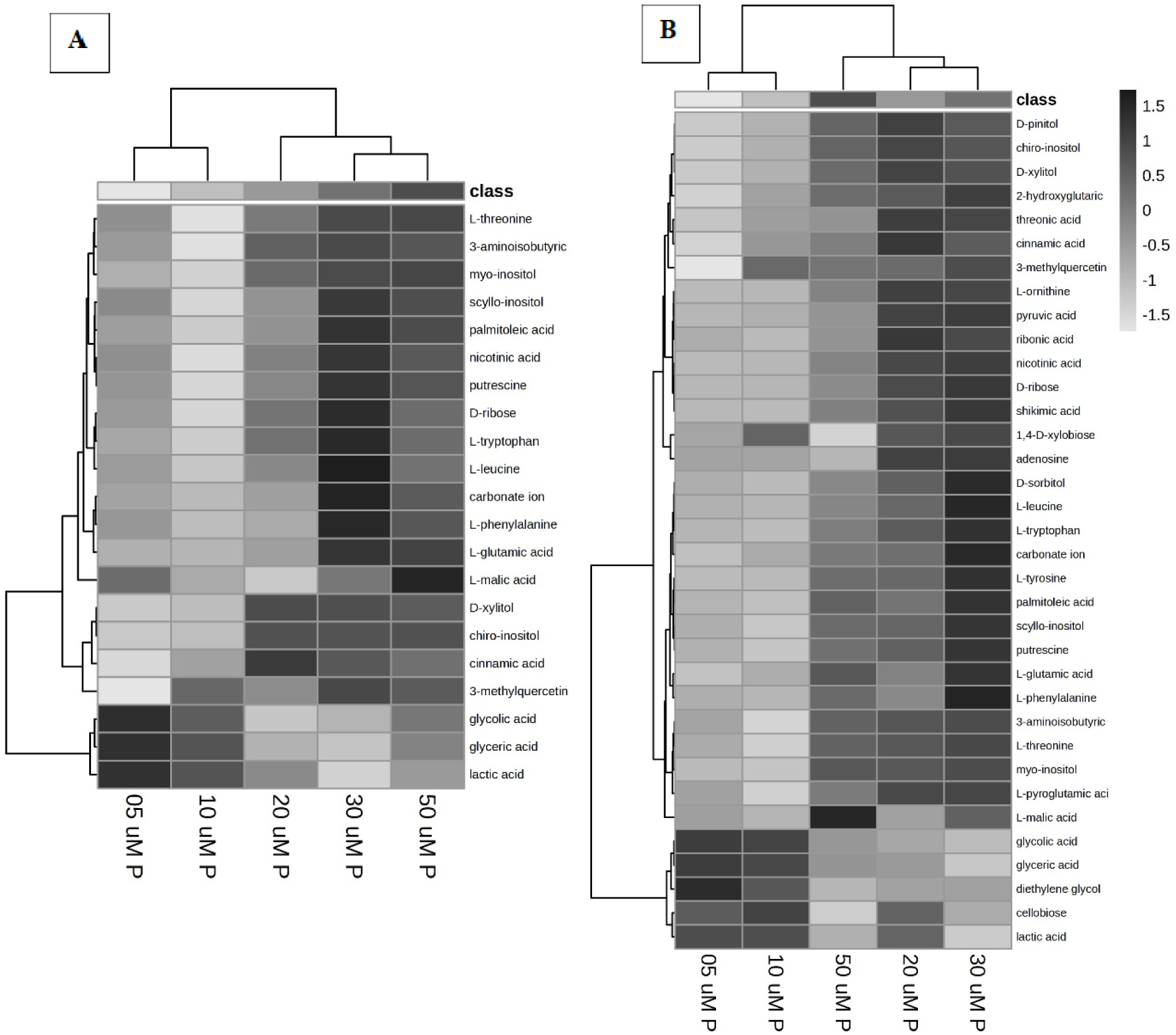
Heatmaps of metabolites showing significant differences among P treatment at flowering (one-way ANOVA, P < 0.05). Hierarchical clustering was performed using Euclidean distance and Ward’s method. Darker shading indicates higher peak areas, whereas lighter shading indicates lower peak areas. (A) Peak areas standardized by root biomass. (B) Peak areas expressed on a per-plant basis.

Overall, these results indicate that the effect of P supply on root exudation was limited at early developmental stages but became pronounced at flowering, when clear compositional shifts were observed between low and higher P treatments.

## Discussion

### Phosphorus effect on shoot and root starts at stem elongation and intensify at flowering

We investigated white lupin development across three phenological stages under a fine P gradient in hydroponics (5, 10, 20, 30 and 50 µM P). Our fine gradient allowed us to capture the transition from P deficiency to sufficiency (Keerthisinghe et al., 1998; O’Sullivan et al., 2021) and revealed that plant responses to P supply were strongly dependent on phenological stage. While previous studies have mostly examined cluster root formation and exudation under extreme P deficiency (Neumann and Römheld, 1999; Massonneau *et al*., 2001; Weisskopf *et al*., 2006; Cheng *et al*., 2014; Müller *et al*., 2015), our approach identified transition response of white lupin to slight increase in P supply. An hydroponic supply of 50 µM P is generally considered sufficient for white lupin growth (Peek *et al*., 2003). However, at flowering, plants supplied with 20-30 µM P achieved similar shoot biomass to those grown at 50 µM P while maintaining greater cluster root formation, suggesting that moderate P limitation optimize root trait development without penalizing shoot growth (Prescott *et al*., 2020; Wang *et al*., 2025).

In contrast, P had little effect during leaf development, despite cotyledons removal intended to induce early P deficiency. This limited early response likely reflected the fine gradient used in our study, and is consistent with finding from O’Sullivan et al., 2022. In contrast, Müller et al., 2015 reported greater root biomass and cluster root production three weeks after plant emergence in a study comparing extreme P deficiencies to very high P supply (500 µM KH_2_PO_4_).

From stem elongation onward, P limitation increasingly affected shoot development. At 5 µM P, shoot biomass, leaf area and specific leaf area were reduced, with differences between P supply intensifying at flowering. Reduced specific leaf area under P limitation likely reflects stronger constraints on cell division and leaf expansion (Freschet *et al*., 2015) than photosynthesis activity (Kavanová *et al*., 2006), potentially leading to an accumulation of excess carbon in leaves and roots (White and Hammond, 2016). In parallel with these changes in shoot traits, low shoot P status is known to control cluster root development (Shane *et al*., 2003; Shane and Lambers, 2005; Li *et al*., 2008) and although cluster root formation was strongly reduced at high P (50 µM P), it was not fully suppressed. This observation aligns with Keerthisinghe et al., 1998, who concluded that cluster root formation and citrate exudation do not necessarily occur at the expense of dry matter production.

Notably, our results indicate that cluster root initiation appeared to cease at flowering and was accompanied by reduced organic acid exudation. To our knowledge, this developmental termination of cluster root initiation in white lupin has not been explicitly reported previously. An explication for this might be an internal P remobilization to sustain flowering instead of relying on external P uptake (Julia *et al*., 2016). Numerous studies have examined the functional differentiation of cluster roots (Massonneau *et al*., 2001) rather than the whole-plant developmental transitions.

Our findings have important implications for strategies aiming to exploit cluster roots and organic acid exudation for mineral dissolution and nutrient acquisition in agroecosystems. Such developmental changes in carbon allocation and nutrient demand are expected to influence root exudate profiles, as exudation is closely linked to plant metabolism.

### Plant phenological stage has a larger effect than P supply on root exudate profiles

The main objective of this study was to disentangle the effect of developmental stage and P supply on white lupin exudate profiles. Our GC-MS-based metabolomics analysis showed that phenological stage exerted a stronger influence than P supply on both the quantity and composition of root exudates. Exudation per root mass was highest during leaf development, followed by stem elongation, and substantially reduced at flowering. This pattern aligns with previous studies showing that actively growing roots release more C into the rhizosphere (Jones *et al*., 2009; Gao *et al*., 2024), and that plants exhibit higher exudation rates per root mass during vegetative growth compared to maturity (Gransee and Wittenmayer, 2000; Aulakh *et al*., 2001; Santangeli *et al*., 2024; Hafner *et al*., 2025). At leaf development, exudates were dominated by compounds originating from central carbon metabolism, including organic acids, amino acids, and sugars, consistent with early-stage exudate profiles in Arabidopsis, pea, rice, and maize (Gransee and Wittenmayer, 2000; Aulakh *et al*., 2001; Chaparro *et al*., 2013; Zhao *et al*., 2021). Although exudation on a per-plant basis was mainly driven by root biomass, 24 compounds were most abundant during leaf development, reflecting the high metabolic activity of young roots. In particular, the elevated exudation of organic acids was linked to the presence of cluster root at this stage (Neumann and Römheld, 2000). By contrast, exudates at later developmental stage shifted towards compounds associated with cell wall turnover, lipid metabolism and secondary pathways. Higher abundance of cell-wall derived compounds may reflect root senescence, particularly cluster roots (Shane and Lambers, 2005).

In parallel with these changes in exudates during plant development, P supply further influenced the exudate composition, although the extent of it depended on the phenological stage. During leaf development, P supply had minimal influence on exudate profiles. During stem elongation, low to moderate P increased the exudation of organic acids, reflecting changes in TCA cycle under P limitation. Increased exudation of amino acids under low P is consistent with the documented increases in amino acid metabolism under P deficiency in white lupin, cotton and soybean (Yan *et al*., 2007; Tawaraya *et al*., 2014; Müller *et al*., 2015). These responses may reflect both N and C being available in surplus when P is limiting (Prescott *et al*., 2020). Higher putrescine exudation under low P treatments might reflect the role of putrescine in stress responses and P remobilization from cell wall (Gill and Tuteja, 2010; Jing *et al*., 2022).

At flowering, exudate profiles of low P (5-10 µM P) were associated with carbohydrates derived from primary C metabolism, cell wall polysaccharide, lipid-related compounds, and phenolics, while moderate to high P showed enrichment in amino acids. The greater release of cell wall–derived sugars under low P may be linked to P foraging strategies involving cell wall P reutilization, as reported in rice (Jing *et al*., 2022), or a result of root cell damage due to P limitation (Niu *et al*., 2012). In parallel, remodelling of cell membrane to replace phospholipids is a well-documented response under phosphate starvation (Russo et al., 2007; White and Hammond 2016). Increased exudation of lipid-related compounds under low P might be a result of this metabolism change. Although flavonoid exudation has previously been reported in white lupin exudates (Weisskopf *et al*., 2006; Cesco *et al*., 2010), these compounds were not detected here, likely reflecting analytical limitations of GC–MS for larger and less volatile secondary metabolites (Ritter *et al*., 2025).

### Implications of shift in exudate profile for rhizosphere-induced ecological benefits

These pronounced shifts in exudate profiles suggest that their effect in the rhizosphere will also be phenological stage-dependent (Oburger et al., 2022). Developmental shifts in exudation are often interpreted to be linked with rhizosphere microbial interactions (van Dam and Bouwmeester, 2016; Vives-Peris *et al*., 2020). Sugars and organic acids released during early growth stages are interpreted to promote microbial recruitment and shape rhizosphere community structure (Zhalnina *et al*., 2018; Canarini *et al*., 2019), whereas aromatic or secondary compounds are more commonly associated with stress or defense response (Badri and Vivanco, 2009). However, these changes may also “simply” reflect metabolic consequence of the plant development and response to environmental changes (Prescott *et al*., 2020).

In addition to microbial community assembly, changes in exudates profiles over plant development is likely to influence other rhizospheric processes such as C and nutrient cycling, and plant nutrient acquisition. Among the most functionally relevant compounds, organic acids play a central role in plant nutrient acquisition strategies (Jones, 1998). In white lupin, organic acid exudation is strongly influenced by the activity of cluster roots. The strong decrease in organic acid exudation as plant developed could impact P acquisition to support flowering. This interpretation is supported by our companion study, in which these exudate cocktails collected at flowering increased Al, Fe, Mg and Ca dissolution in three contrasting soil types, but had only limited effect on P dissolution (Pollet *et al*., 2026). Together, these results suggest that exudates from younger plants would likely induce substantially greater P mobilization.

Higher organic acid exudation from younger plants may also have consequences for soil C dynamics. Organic acids such as oxalic acid can disrupt organo-mineral association and destabilize previously stable soil organic matter (Keiluweit *et al*., 2015). In parallel, the greater exudation of readily metabolizable sugars may stimulate microbial activity and growth, potentially enhancing microbially mediated mechanisms, such as rhizosphere priming leading to substantial C degradation (Li *et al*., 2021; Zhang *et al*., 2022). Furthermore, the release of non-polar and hydrophobic content such as lipid may interact differently with rhizospheric components compared with neutral and polar compounds (e.g. amino acids) or charged compounds (e.g. organic acid) (Bölscher et al., 2025), with potential consequence for organo-mineral associations, nutrient cycling and microbial community composition (Couvillion et al., 2025).

Some limitation of this study should nevertheless be acknowledged. Because the hydroponic system was not sterile, part of the detected metabolites may have originated from microorganisms rather than roots alone. Simultaneously, the lower root-to-sampling solution volume ratio at leaf development (1.2 g□¹ root dry weight L□¹ sampling solution) in comparison to flowering (4.6 g□¹ root L□¹ solution) might have slightly overestimated exudation at early stage (Otxandorena-Ieregi *et al*., 2024). Finally, exudation patterns observed here under hydroponic conditions may not fully represent those occurring under field conditions. as substantial differences in exudates composition between plants grown in the hydroponics system and those grown in soil exist (Williams *et al*., 2021; Heuermann *et al*., 2023).

Taken together, our findings highlight the strongly dynamic nature of root exudation across plant development and to some extent P limitation. Our study identified more than 100 metabolites in white lupin exudates, yet the ecological significance of many of these compounds remains poorly understood. Beyond well-characterized organic acids, it is still unclear which metabolic pathways produce many of these metabolites, how they are regulated by plant development and P availability, and what their fate and function are in the rhizosphere. A better understanding of these temporal changes is essential for linking plant phenology with rhizosphere functioning and for improving the use of plant–soil interactions to support more sustainable agroecosystems.

## Supplementary data

Supplementary material 1: Summary of plant parameters and results from two-way ANOVA testing the effect of phenological stage, P treatment and their interaction.

Supplementary material 2: Summary of quantified exudate compounds by GC-MS and results from two-way ANOVA testing the effect of phenological stage, P treatment and their interaction.

Supplementary material 3: List of metabolites in white lupin exudates and their chemical classes.

Supplementary material 4: Results of the PCA on peak area expressed on a per-plant basis across the three phenological stages.

## Acknowledgements

We would like to thank colleagues from the University of British Columbia, particularly Fiona Farag and Mary Clare Kendrick for their help in experimental set up. We gratefully acknowledge Hatem Rouached for his constructive pre-submission feedback.

## Author contributions

JT-C and SP: conceptualization; SP, JT-C and GL: methodology; YM-K, KT: metabolomic analysis; SP, JT-C and GL investigation; SP: data analysis; SP: writing - original draft; SP, JT-C, GL, TK, CP : writing - review & editing; SP: visualization; JT-C, GL, CP and TK: supervision; JT-C: funding acquisition

## Conflict of interest

No conflict of interest declared

## Funding

Sasha Pollet is supported by the Wallonia-Brussels International scholarship. This project [GIR003] is supported by the Genomic Innovation for Regenerative Agriculture, Food and Fisheries (GIRAFF) Program, which is funded by the Governments of British Columbia and Canada and delivered by Genome British Columbia and the Investment Agriculture Foundation of BC A portion of this research was performed under the Large-Scale Research program (https://doi.org/10.46936/lser.proj.2022.60372/60008596) at the Environmental Molecular Sciences Laboratory located in Pacific Northwest National Laboratory (PNNL), a Department of Energy (DOE) Office of Science User Facility sponsored by the Biological and Environmental Research (BER) program. Battelle operates PNNL for DOE under Contract No. DE-AC05-76RL01830.

## Notes

### Competing Interest Statement

The authors have declared no competing interest.

## References

Aulakh MS, Wassmann R, Bueno C, Kreuzwieser J, Rennenberg H. 2001. Characterization of root exudates at different growth stages of ten rice (Oryza sativa L.) cultivars. Plant Biology 3, 139–148.

Badri D V., Loyola-Vargas VM, Broeckling CD, et al. 2008. Altered profile of secondary metabolites in the root exudates of arabidopsis ATP-binding cassette transporter mutants. Plant Physiology 146, 762–771.

Badri D V., Vivanco JM. 2009. Regulation and function of root exudates. Plant, cell & environment 32, 666–681.

Brown RW, Reay MK, Centler F, Chadwick DR, Bull ID, McDonald JE, Evershed RP, Jones DL. 2024. Soil metabolomics - current challenges and future perspectives. Soil Biology and Biochemistry 193, 109382.

Bunn RA, Corrêa A, Joshi J, Kaiser C, Lekberg Y, Prescott CE, Sala A, Karst J. 2024. What determines transfer of carbon from plants to mycorrhizal fungi? New Phytologist 244, 1199–1215.

Canarini A, Kaiser C, Merchant A, Richter A, Wanek W. 2019. Root exudation of primary metabolites: Mechanisms and their roles in plant responses to environmental stimuli. Frontiers in Plant Science 10, 157.

Carvalhais LC, Dennis PG, Fedoseyenko D, Hajirezaei MR, Borriss R, Von Wirén N. 2011. Root exudation of sugars, amino acids, and organic acids by maize as affected by nitrogen, phosphorus, potassium, and iron deficiency. Journal of Plant Nutrition and Soil Science 174, 3–11.

Cesco S, Neumann G, Tomasi N, Pinton R, Weisskopf L. 2010. Release of plant-borne flavonoids into the rhizosphere and their role in plant nutrition. Plant and Soil 329, 1–25.

Chaparro JM, Badri D V., Bakker MG, Sugiyama A, Manter DK, Vivanco JM. 2013. Root Exudation of Phytochemicals in Arabidopsis Follows Specific Patterns That Are Developmentally Programmed and Correlate with Soil Microbial Functions. PLOS ONE 8, e55731.

Cheng L, Tang X, Vance CP, White PJ, Zhang F, Shen J. 2014. Interactions between light intensity and phosphorus nutrition affect the phosphate-mining capacity of white lupin (Lupinus albus L.). Journal of Experimental Botany 65, 2995–3003.

van Dam NM, Bouwmeester HJ. 2016. Metabolomics in the Rhizosphere: Tapping into Belowground Chemical Communication. Trends in Plant Science 21, 256–265.

Dietz S, Herz K, Gorzolka K, Jandt U, Bruelheide H, Scheel D. 2020. Root exudate composition of grass and forb species in natural grasslands. Scientific Reports 2020 10:1 10, 1–15.

Feng H, Fu R, Luo J, et al. 2023. Listening to plant’s Esperanto via root exudates: reprogramming the functional expression of plant growth-promoting rhizobacteria. New Phytologist 239, 2307–2319.

Freschet GT, Swart EM, Cornelissen JHC. 2015. Integrated plant phenotypic responses to contrasting above- and below-ground resources: Key roles of specific leaf area and root mass fraction. New Phytologist 206, 1247–1260.

Gao Y, Wang H, Yang F, Dai X, Meng S, Hu M, Kou L, Fu X. 2024. Relationships between root exudation and root morphological and architectural traits vary with growing season. Tree Physiology 44.

Gill SS, Tuteja N. 2010. Polyamines and abiotic stress tolerance in plants. Plant Signaling and Behavior 5, 26–33.

Gransee A, Wittenmayer L. 2000. Qualitative and quantitative analysis of water-soluble root exudates in relation to plant species and development. Journal of Plant Nutrition and Soil Science 163, 381–385.

Hafner BD, Pietz O, King WL, Scharfetter JB, Bauerle TL. 2025. Early developmental shifts in root exudation profiles of five Zea mays L. genotypes. Plant Science 354.

Han M, Chen Y, Sun L, Yu M, Li R, Li S, Su J, Zhu B. 2023. Linking rhizosphere soil microbial activity and plant resource acquisition strategy. Journal of Ecology 111, 875–888.

Hartmann H, Bahn M, Carbone M, Richardson AD. 2020. Plant carbon allocation in a changing world – challenges and progress: introduction to a Virtual Issue on carbon allocation: Introduction to a virtual issue on carbon allocation. New Phytologist 227, 981–988.

Heuermann D, Döll S, Schweneker D, Feuerstein U, Gentsch N, von Wirén N. 2023. Distinct metabolite classes in root exudates are indicative for field- or hydroponically-grown cover crops. Frontiers in Plant Science 14, 1122285.

Hinsinger P. 2001. Bioavailability of soil inorganic P in the rhizosphere as affected by root-induced chemical changes: A review. Plant and Soil 237, 173–195.

Jing HK, Wu Q, Huang J, Yang XZ, Tao Y, Shen RF, Zhu XF. 2022. Putrescine is involved in root cell wall phosphorus remobilization in a nitric oxide dependent manner. Plant Science 316, 111169.

Jones DL. 1998. Organic acids in the rhizosphere-a critical review. Plant and Soil 205, 25–44.

Jones DL, Hodge A, Kuzyakov Y. 2004. Plant and mycorrhizal regulation of rhizodeposition. New Phytologist 163, 459–480.

Jones DL, Nguyen C, Finlay RD. 2009. Carbon flow in the rhizosphere: Carbon trading at the soil-root interface. Plant and Soil 321, 5–33.

Kanehisa M, Furumichi M, Sato Y, Matsuura Y, Ishiguro-Watanabe M. 2025. KEGG: biological systems database as a model of the real world. Nucleic acids research 53, D672–D677.

Kassambara A. 2023. rstatix: Pipe-Friendly Framework for Basic Statistical Tests. in press.

Kavanová M, Lattanzi FA, Grimoldi AA, Schnyder H. 2006. Phosphorus Deficiency Decreases Cell Division and Elongation in Grass Leaves. Plant Physiology 141, 766.

Keerthisinghe G, Hocking PJ, Ryan PR, Delhaize E. 1998. Effect of phosphorus supply on the formation and function of proteoid roots of white lupin (Lupinus albus L.). Plant, Cell and Environment 21, 467–478.

Keiluweit M, Bougoure JJ, Nico PS, Pett-Ridge J, Weber PK, Kleber M. 2015. Mineral protection of soil carbon counteracted by root exudates. Nature Climate Change 5, 588–595.

Kim S, Chen J, Cheng T, et al. 2025. PubChem 2025 update. Nucleic Acids Research 53, D1516–D1525.

Kind T, Wohlgemuth G, Lee DY, Lu Y, Palazoglu M, Shahbaz S, Fiehn O. 2009. FiehnLib: Mass spectral and retention index libraries for metabolomics based on quadrupole and time-of-flight gas chromatography/mass spectrometry. Analytical Chemistry 81, 10038–10048.

Li H, Testerink C, Zhang Y. 2021. How roots and shoots communicate through stressful times. Trends in Plant Science 26, 940–952.

Liu X, Heinzle J, Tian Y, Salas E, Borken W, Schindlbacher A, Wanek W. 2025. Primary metabolites in root exudates are not affected by long-term soil warming in a temperate forest. Functional Ecology 40, 417–432.

Lobet G, Pagès L, Draye X. 2011. A novel image-analysis toolbox enabling quantitative analysis of root system architecture. Plant Physiology 157, 29–39.

Lynch JP, Brown KM. 2016. Root strategies for phosphorus acquisition. In: White P, Hammond J, eds. The Ecophysiology of Plant-Phosphorus interaction. 83–116.

Lynch JM, Whipps JM. 1990. Substrate flow in the rhizosphere. Plant and Soil 129, 1–10.

Ma W, Tang S, Dengzeng Z, Zhang D, Zhang T, Ma X. 2022. Root exudates contribute to belowground ecosystem hotspots: A review. Frontiers in Microbiology 13, 937940.

Massonneau A, Langlade N, Léon S, Smutny J, Vogt E, Neumann G, Martinoia E. 2001. Metabolic changes associated with cluster root development in white lupin (Lupinus albus L.): Relationship between organic acid excretion, sucrose metabolism and energy status. Planta 213, 534–542.

McCulloch LA, Taylor BN, Wurzburger N, Prescott CE. 2026. Rethinking symbiotic nitrogen fixation: Could surplus carbon drive unexpected patterns of resource allocation? Plant and Soil doi: 10.1007/s11104-026-08391-0.

McLaughlin S, Zhalnina K, Kosina S, Northen TR, Sasse J. 2023. The core metabolome and root exudation dynamics of three phylogenetically distinct plant species. Nature Communications 14, 1–13.

Müller J, Gödde V, Niehaus K, Zörb C. 2015. Metabolic adaptations of white lupin roots and shoots under phosphorus deficiency. Frontiers in Plant Science 6, 1–10.

Münch E. 1930. Die stoffbewegungen in der pflanze. Stoffbewegungen in der pflanze in press.

Neumann G, Römheld V. 1999. Root excretion of carboxylic acids and protons in phosphorus-deficient plants. Plant and Soil 211, 121–130.

Neumann G, Römheld V. 2000. The Release of Root Exudates as Affected by the Plant Physiological Status. In: Willig S, Varanini Z, Nannipieri P, eds. The Rhizosphere: Biochemistry and Organic Substances at the Soil-Plant Interface.

Neumann G, Römheld V. 2011. Rhizosphere Chemistry in Relation to Plant Nutrition. Marschner’s Mineral Nutrition of Higher Plants: Third Edition. Elsevier Inc., 347–368.

New South Wales. Department of Industry and Investment. 2011. Lupin growth & development / NSW Government Industry & Investment. (J Edwards and J Walker, Eds.). State of New South Wales: NSW Dept. of Industry & Investment.

Niu YF, Chai RS, Jin GL, Wang H, Tang CX, Zhang YS. 2012. Responses of root architecture development to low phosphorus availability: a review. Annals of Botany 112, 391.

Oburger E, Jones DL. 2018. Sampling root exudates – Mission impossible? Rhizosphere 6, 116–133.

Oburger E, Schmidt H, Staudinger C. 2022. Harnessing belowground processes for sustainable intensification of agricultural systems. Plant and Soil 2022 478:1 478, 177–209.

O’Sullivan JB, Jin J, Tang C. 2022. White lupin (Lupinus albus L.) exposed to elevated atmospheric CO2 requires additional phosphorus for N2 fixation. Plant and Soil 476, 477–490.

O’Sullivan JB, Plozza T, Stefanelli D, Jin J, Tang C. 2021. Elevated CO2 and phosphorus deficiency interactively enhance root exudation in Lupinus albus L. Plant and Soil 465, 229–243.

Otxandorena-Ieregi U, Santangeli M, Aleksza D, Hann S, Oburger E. 2024. Fine-tuning root exudation sampling procedures– evaluating the effect of sampling solution volume and the suitability of Micropur as microbial activity inhibitor. Plant and Soil 2024 504:1 504, 415–433.

Pang Z, Lu Y, Zhou G, et al. 2024. MetaboAnalyst 6.0: towards a unified platform for metabolomics data processing, analysis and interpretation. Nucleic Acids Research 52, W398–W406.

Pantigoso HA, Manter DK, Vivanco JM. 2020a. Differential Effects of Phosphorus Fertilization on Plant Uptake and Rhizosphere Microbiome of Cultivated and Non-cultivated Potatoes. Microbial Ecology 2020 80:1 80, 169–180.

Pantigoso HA, Yuan J, He Y, Guo Q, Vollmer C, Vivancoid JM. 2020b. Role of root exudates on assimilation of phosphorus in young and old Arabidopsis thaliana plants. doi: 10.1371/journal.pone.0234216.

Pausch J, Kuzyakov Y. 2018. Carbon input by roots into the soil: Quantification of rhizodeposition from root to ecosystem scale. Global Change Biology 24, 1–12.

Peek CS, Robson AD, Kuo J. 2003. The formation, morphology and anatomy of cluster root of Lupinus albus L. as dependent on soil type and phosphorus supply. Plant and Soil 248, 237–246.

Plaxton WC, Tran HT. 2011. Metabolic Adaptations of Phosphate-Starved Plants. Plant Physiology 156, 1006.

Pollet S, Cornelis J-T, Knipfer T, Prescott C, Tate K, Kim Y-M, Lobet G. 2026. How exudates production along a phosphorus gradient influences mineral dissolution across contrasting soil development stages. Plant and Soil, 1–22.

Prescott CE, Grayston SJ, Helmisaari HS, Kaštovská E, Körner C, Lambers H, Meier IC, Millard P, Ostonen I. 2020. Surplus Carbon Drives Allocation and Plant–Soil Interactions. Trends in Ecology and Evolution 35, 1110–1118.

Prescott CE, Rui Y, Cotrufo MF, Grayston SJ. 2021. Managing plant surplus carbon to generate soil organic matter in regenerative agriculture. Journal of Soil and Water Conservation 76, 99A–104A.

R Core Team. 2022. R: A language and environment for statistical computing. R Foundation for Statistical Computing in press.

Ren P, Li P, Zhou X, Liu Z, Tang J, Zhang C, Zou Z, Li T, Peng C. 2024. Shifts in Plant Phenology Significantly Affect the Carbon Allocation in Different Plant Organs. Ecology Letters 27.

Ritter A, Phillip ·, Becker J, Möller K, Granse D, Jensen K, Meier IC, Harihar ·, Subrahmaniam J, Becker PJ. 2025. Targeting the untargeted: Uncovering the chemical complexity of root exudates. Plant and Soil 2025, 1–16.

Robert CAM, Himmighofen P, Mclaughlin S, Cofer TM, Khan SA, Siffert A, Sasse J. 2026. Environmental and Biological Drivers of Root Exudation. Annual Review of Plant Biology Downloaded from www.annualreviews.org. Guest 48, 2.

Salem MA, Wang JY, Al-Babili S. 2022. Metabolomics of plant root exudates: From sample preparation to data analysis. Frontiers Media S.A.

Santangeli M, Steininger-Mairinger T, Vetterlein D, Hann S, Oburger E. 2024. Maize (Zea mays L.) root exudation profiles change in quality and quantity during plant development – A field study. Plant Science 338, 111896.

Seethepalli A, York LM. 2020. RhizoVision Explorer - Interactive software for generalized root image analysis designed for everyone. doi: 10.5281/zenodo.4095629.

Shane MW, Lambers H. 2005. Cluster roots: A curiosity in context. Plant and Soil 274, 101–125.

Shane MW, De Vos M, De Roock S, Lambers H. 2003. Shoot P status regulates cluster-root growth and citrate exudation in Lupinus albus grown with a divided root system. Plant, Cell and Environment 26, 265–273.

Sumner LW, Amberg A, Barrett D, et al. 2007. Proposed minimum reporting standards for chemical analysis. Metabolomics 3, 211–221.

Tawaraya K, Horie R, Shinano T, Wagatsuma T, Saito K, Oikawa A. 2014. Metabolite profiling of soybean root exudates under phosphorus deficiency. Soil Science and Plant Nutrition 60, 679–694.

Vengavasi K, Pandey R. 2018. Root exudation potential in contrasting soybean genotypes in response to low soil phosphorus availability is determined by photo-biochemical processes. Plant Physiology and Biochemistry 124, 1–9.

Vitousek PM, Porder S, Houlton BZ, Chadwick OA. 2010. Terrestrial phosphorus limitation: mechanisms, implications, and nitrogen–phosphorus interactions. Ecological Applications 20, 5–15.

Vives-Peris V, de Ollas C, Gómez-Cadenas A, Pérez-Clemente RM. 2020. Root exudates: from plant to rhizosphere and beyond. Plant Cell Reports 39, 3–17.

Wang C, Pollet S, Howell K, Cornelis J-T. 2025. Placing cropping systems under suboptimal phosphorus conditions promotes plant nutrient acquisition and microbial carbon supply without compromising biomass. Soil Biology and Biochemistry, 109753.

Wasaki J, Rothe A, Kania A, Neumann G, Römheld V, Shinano T, Osaki M, Kandeler E. 2005. Root Exudation, Phosphorus Acquisition, and Microbial Diversity in the Rhizosphere of White Lupine as Affected by Phosphorus Supply and Atmospheric Carbon Dioxide Concentration. Journal of Environmental Quality 34, 2157–2166.

Weisskopf L, Tomasi N, Santelia D, Martinoia E, Langlade NB, Tabacchi R, Abou-Mansour E. 2006. Isoflavonoid exudation from white lupin roots is influenced by phosphate supply, root type and cluster-root stage. New Phytologist 171, 657–668.

White P, Hammond J. 2016. Phosphorus nutrition of terrestrial plants. The Ecophysiology of Plant-Phosphorus interaction. 51–81.

Williams A, Langridge H, Straathof AL, Fox G, Muhammadali H, Hollywood KA, Xu Y, Goodacre R, de Vries FT. 2021. Comparing root exudate collection techniques: An improved hybrid method. Soil Biology and Biochemistry 161.

Williams A, de Vries FT. 2020. Plant root exudation under drought: implications for ecosystem functioning. New Phytologist 225, 1899–1905.

Xu L, Naylor D, Dong Z, et al. 2018. Drought delays development of the sorghum root microbiome and enriches for monoderm bacteria. Proceedings of the National Academy of Sciences of the United States of America 115, E4284–E4293.

Yan WD, Shi WM, Li BH, Zhang M. 2007. Overexpression of a Foreign Bt Gene in Cotton Affects the Low-Molecular-Weight Components in Root Exudates. Pedosphere 17, 324–330.

Zhalnina K, Louie KB, Hao Z, et al. 2018. Dynamic root exudate chemistry and microbial substrate preferences drive patterns in rhizosphere microbial community assembly. Nature Microbiology 2018 3:4 3, 470–480.

Zhang D, Zhang Y, Zhao Z, Xu S, Cai S, Zhu H, Rengel Z, Kuzyakov Y. 2022. Carbon–Phosphorus Coupling Governs Microbial Effects on Nutrient Acquisition Strategies by Four Crops. Frontiers in Plant Science 13, 924154.

Zhao M, Zhao J, Yuan J, Hale L, Wen T, Huang Q, Vivanco JM, Zhou J, Kowalchuk GA, Shen Q. 2021. Root exudates drive soil-microbe-nutrient feedbacks in response to plant growth. Plant, Cell & Environment 44, 613–628.

